# Skeletal maturity predicts cognitive abilities in human adolescents

**DOI:** 10.1101/2021.05.02.442351

**Authors:** Ilona Kovács, Kristóf Kovács, Patrícia Gerván, Katinka Utczás, Gyöngyi Oláh, Zsófia Tróznai, Andrea Berencsi, Hanna Szakács, Ferenc Gombos

**Author notes:** **corresponding author**: Ilona Kovács, Laboratory for Psychological Research, Pázmány Péter Catholic University, 1 Mikszáth sq., Budapest, 1088 Hungary.

## Abstract

Adolescent human development is not only shaped by the mere passing of time and accumulating experience, but it also depends on pubertal timing and the cascade of maturational processes orchestrated by gonadal hormones. Although individual variability in puberty onset confounds adolescent studies, it has not been efficiently controlled for. Here we introduce ultrasonic bone age assessment to estimate biological maturity and disentangle the independent effects of chronological and biological age on adolescent cognitive abilities. Comparing cognitive performance of participants with different skeletal maturity we uncover the striking impact of biological age on both IQ and specific abilities. We find that biological age has a selective effect on abilities: more mature individuals within the same age group have higher working memory capacity and processing speed, while those with higher chronological age have better verbal abilities, independently of their maturity. Bone age appears to be a surprisingly strong predictor of cognitive abilities, and it seems that a teen’s IQ is determined by biological age.

## Introduction

Entering a classroom full of first year high-school students is a peculiar experience as we meet pupils who are of the same age, but extremely diverse in terms of stature, behavioural features, social and emotional outlook and even cognitive abilities. This is because adolescent growth and development are not only shaped by the mere passing of time, but they also depend on the timing of puberty onset, which has a large variability across individuals^1–3^. After the onset of puberty, musculoskeletal, reproductive and neurodevelopmental systems of a child start to transform into those of an adult during a period that is extremely malleable but also vulnerable. Our current knowledge is limited regarding the correspondence between physical and psychological maturity in adolescence. Nevertheless, we can hypothesize that apparent changes in physique reflecting hormonal progress are accompanied by the structural remodelling of the brain^4–7^ and followed by functional advancement^8–10^.

Although the variability in the onset of puberty results in discrepancies with respect to developmental trajectories^1,11–13^, such variability has not been very efficiently controlled for in adolescent research^1,4,12^. Additionally, the methods of assessment are mostly outdated, and the dissociation of chronological and biological age has not been addressed clearly. The gold-standard has been the Tanner Scale^14–16^, which is based on bodily features such as breast and testicle sizes, and other secondary sex characteristics visually assessed by a paediatrician. This method is very subjective, it might be unsettling for the subject, and it does not seem to represent contemporary nutritional conditions and secular trends of human growth^3,13,15,16^. The Tanner Scale is based on a post-war longitudinal study carried out in the Harpenden children’s home near London, between 1949 and 1971^14,16^. The Harpenden Growth Study^14,18^ involved neglected, many times orphaned children whose participation would not only raise a number of ethical issues today^16^, but it is highly questionable whether the Tanner Scale can serve as a reliable and valid standard of developmental stages in the present^15,16^.

We believe that more objective indicators of biological age, such as gonadal hormone levels^19^, metabolic markers^20^, genetic indicators^21^, skeletal^22^ or even dental^23^ maturity assessments might be better alternatives to replace Tanner staging in developmental research and serve as biomarkers of pubertal onset and progress. The challenge is to identify an indicator that is correlated with adolescent hormonal development and, at the same time, is also sensitive enough to identify teenagers who are either advanced or delayed compared to their peers. Expense and burden to the subject also needs to be considered both in large population surveys and in laboratory studies.

Bone age has been widely used in paediatric practice^24,25^, and it is generally based on hand and wrist X-ray radiographs assessing the size and geometry of the epiphysis and fusion of the epiphysis and diaphysis. It has been shown that pubertal hormone levels are correlated more with skeletal system development and bone age than with chronological age^26–28^. This raises the possibility that bone age might be a useful proxy of pubertal maturity. The prospect of a bone age-based pubertal maturity assessment paradigm for research is enhanced by a non-invasive ultrasonic version of this method that scans the ossification of the wrist region to estimate bone age^29^. We have shown that ultrasonic bone age estimations of the wrist very strongly correlate with X-ray estimations^30^ (r>0.95), and therefore provide a promising tool to replace subjective, invasive, risky or costly assessments of maturity.

In this study, biological age (BA) of participants is assessed by evaluating the acoustic parameters of the left wrist, establishing bone age for each participant (Methods/Procedure). In order to estimate the independent effects of BA and chronological age (CA) on development we introduced the paradigm presented in Figure 1. Preselected participants along the Maturity axis of Fig. 1a have the same CA and different BAs, therefore, any differences in their cognitive performance will be accounted for exclusively by BA without the influence of CA. And vice versa, performance of participants with different CAs along the Experience axis will only be determined by CA as there is no variability in BA. Aiming at this clear-cut dissociation, we identified cohorts of adolescent participants with average (BA=CA), advanced (BA>CA) and delayed pace of skeletal maturation (BA<CA, Fig. 1b) reflecting their pubertal maturity levels. This design allows for predictions with respect to developmental scenarios where the amount of experience (CA) or the level of maturity (BA) might have different contributions to the studied psychological function (Fig. 1c).

**Fig. 1:**
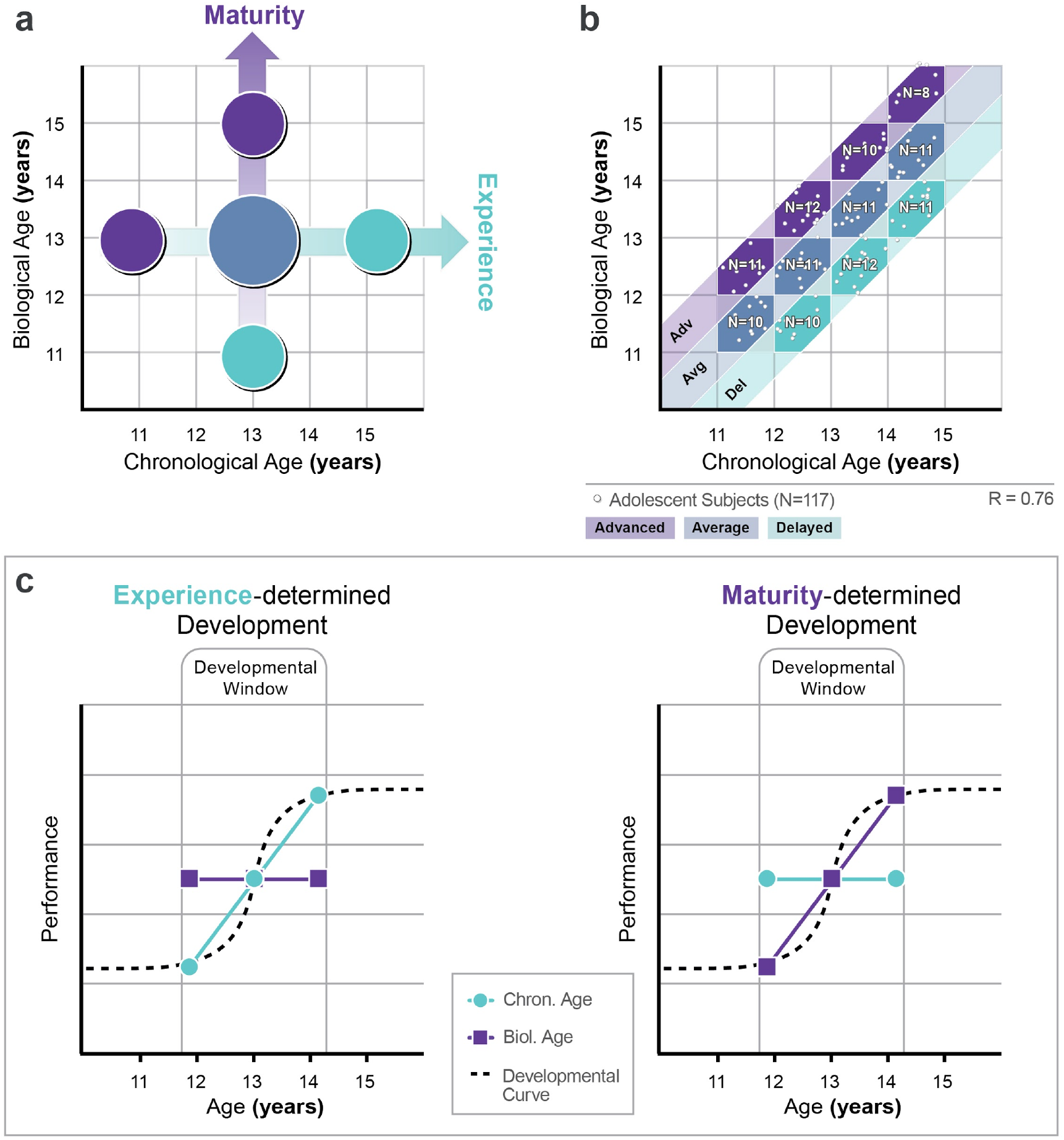
Dissociating the effects of experience and maturation in adolescent development. **a**, Comparing cognitive performance of participants with different biological age (BA) within the same chronological age-group can reveal the impact of biological age (vertical ‘Maturity’ axis) independently of chronological age (CA), while the comparison of different CA participants with the same maturity level reveals the impact of CA independently of maturation (horizontal “Experience” axis). **b**, In order to dissociate the impact of BA and CA we identified cohorts of participants with advanced, average and delayed maturity using bone age assessment within 11 and 15 years of CA. Each one-year-wide hexagonal bin consisted of 8 to 12 subjects (white dots within the bins). **c**, Predictions: Vertical axes indicate development in any cognitive domain. Horizontal axes express age (CA or BA). Important developmental events occur within a “Developmental window” where performance radically changes approaching adult levels. In the extreme case of experience-determined development, we expect to see a significant improvement of performance between different CA groups (matched for BA), while performance will remain the same across different BA groups (matched for CA). In the other extreme case where development is solely maturity-dependent, we expect improvement across different BA groups, and the lack of change across different CA groups.

Predominantly experience-determined development will result in a steep developmental function comparing subjects along the Experience axis of Fig. 1a, while more maturity-determined development will generate a steep function comparing participants along the Maturity axis. It is important to note that the “developmental window” within which a particular psychological function significantly improves might shift across different age-bands depending on the timing of the maturation of the underlying cortical structures. To determine narrower age-bands where the impact of BA/CA might be prominent we generated a number of distinct BA-CA bins shown in Fig. 1b (for a more detailed definition of BA-CA bins see Methods/Participants).

The paradigm introduced in Fig 1. provides a tool to study development in any particular domain. We chose to assess the relationship between cognitive development and skeletal maturity. There has been very limited research in this area, and it is mostly related to extreme skeletal maturational lags that can negatively impact intellectual development^31,32^. Our purpose is to assess the relationship between intellectual development and pubertal maturity within the typical range of developmental parameters. In the lack of extended previous research, we opted for the mapping of different broad cognitive abilities measured by the Wechsler Intelligence Scale for Children.

Traditionally, age effects have been among the primary means towards fractionating human cognitive abilities. In particular, the distinction between fluid and crystallized intelligence, the forerunner of one of the most respected psychological theories on the structure of cognitive abilities, the Cattell-Horn-Carroll (CHC) model^33^, was to a large extent based on the finding that fluid reasoning (the ability to solve novel problems) show developmental patterns very different from crystallized intelligence (the ability to use already acquired skills and knowledge).

Different trajectories of cognitive development provide another means towards dissociating aspects of cognition. For instance, executive functions keep maturing well into the late teenage years, perhaps even until the early twenties^9,10,34^. Performance on different ability tests indeed have very different peaks in life: while tests of memory or processing speed peak as early as the end of teenage years, performance on verbal tests such as those measuring vocabulary or comprehension peaks at near 50 years of age, indicating that experience matters much more for verbal comprehension than for other abilities^35^.

To explore the independent and potentially selective effects of maturation and experience on the cognitive development of a preselected cohort of adolescents (Fig. 1b), we administered 11 subtests from the Hungarian adaptation of the Wechsler Intelligence Scale for Children IV - WISC-IV^36^. According to the hypotheses presented in Fig. 1c, we expected to see a pattern in the results where certain cognitive abilities are more determined by BA, and others by CA. In particular, according to previous results^35,37^ and the theory of fluid and crystallized intelligence^33,38^, we expected verbal abilities to be more influenced by chronological age (probably reflecting the amount of schooling) than non-verbal abilities.

Indeed, our most striking finding is that while verbal abilities are independently influenced by experience (CA has a significant effect when BA is partialled out), the opposite is the case for working memory and processing speed (BA has a significant effect when CA is controlled for). That is, in adolescents of the exact same age, those who are more mature have higher memory capacity and mental speed, while among adolescents at the same level of maturation, those with a higher CA have better verbal abilities.

## Results

Following the assessment of their bone age, participants (11- to 15-year-old, N=117, see Methods and Fig. 1b) were administered 11 WISC-IV subtests (see Methods/Procedure). We only tested female participants in order to control for the different maturational trajectories resulting from a different hormonal composition between the two sexes^13,39–41^. To control for the impact of different socio-economic and sociocultural backgrounds^9,40,42^, we restricted the participant pool to only include pupils arriving from top-level high schools (see Methods/Participants).

Since we purported to study the effect of age-related factors on test performance across age groups, age-standardized scores were not suitable. Therefore, we used unstandardized raw scores, in percentage, for each subtest. Overall performance was calculated by averaging the 11 subtests. We also calculated scores for each of the broad abilities reflected by the indices of the WISC-IV (Verbal Comprehension, Perceptual Reasoning, Working Memory, Processing Speed) by averaging the subtest scores corresponding to each. See Supplementary Tables 1 and 2 for descriptive statistics and basic correlations.

### Overall cognitive performance is determined both by biological and chronological age

In order to obtain a comprehensive perspective on mental abilities first, we calculated and averaged the overall performance of participants for each BA and CA defined one-year bin of Fig. 1b. Overall performance on WISC-IV for each bin is plotted in Fig. 2a. Unsurprisingly, there is increasing performance throughout the entire CA range as adolescents’ cognitive development takes place (remember, the performance measure reflects absolute performance, unlike age-standardised scores such as IQ). A more fascinating trend, however, is the increasing performance as a function of BA. It appears that participants with the same CA but with more advanced bone maturity achieve higher scores on the IQ test!

**Fig. 2:**
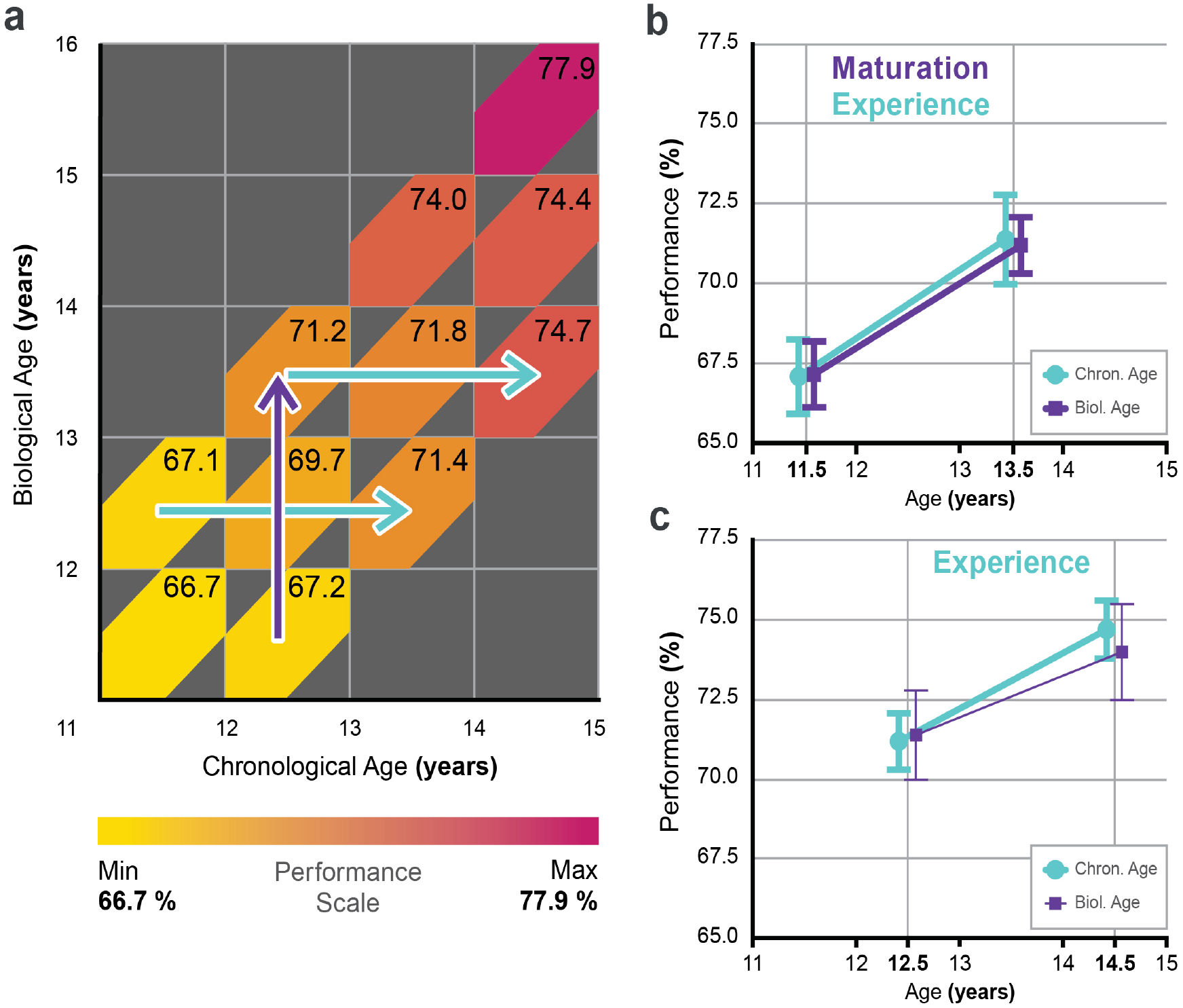
WISC-IV overall performance as a function of biological and chronological age. **a**, Average WISC-IV overall performance (based on raw scores) was calculated for each bin as a % of possible maximum performance described in Fig.1b and expressed on a color scale (Performance scale). Higher performance is indicated by darker shades of red. Numbers within each colored bin express performance in % of possible maximum performance. → indicates bins where a significant biological age effect is present. → indicates bins where a significant chronological age effect is present. **b**, Detailed performance within the 11-14 years age-range where significant BA and CA effects according to one-way ANOVA are present (significant developmental changes are represented by bold lines). Both experience and maturation significantly determine performance. **c**, Detailed performance within the 12-15 years age-range where significant chronological age-effect is present. Performance is determined by experience. Error bars show 1SE.

Pearson correlation was calculated between overall WISC performance and BA/CA within each one-year wide BA or CA group. The correlations were Holm-Bonferroni corrected for multiple comparisons. We used one-way ANOVA to look for significant differences between the lower and upper bins within the particular BA/CA age-band with significant correlations. Coloured arrows in Fig 2a indicate those BA/CA age-bands where significant differences were found.

The two axes in Fig. 1a provide a clear dissociation of maturity (BA) and experience (CA) effects. Since the developmental window (Fig. 1c) for cognitive development might shift across different age ranges depending on timing of the maturation of the underlying cortical structures, it is important to analyse the results within narrower age-ranges and identify the age where the impact of CA/BA is most prominent. Fig. 2b presents detailed analysis within a developmental window of 11 to 14 years of CA/BA with CA comparisons for BA being constant at 12.5, and BA comparisons for CA being constant at 12.5. Apparently, both CA (67.1 % ± 1.2 SE vs. 71.4 % ± 1.4 SE; p = 0.029) and BA (67.2 % ± 1.0 SE vs. 71.2 % ± 0.9 SE; p = 0.007) significantly determine cognitive performance within this CA/BA range. In terms of the predictions of Fig. 1c, development within this window is determined both by experience and maturation. Moving to the next developmental window of 12 to 15 years of CA/BA (Fig. 2c) there seems to be a similarly strong CA effect (71.2 % ± 0.9 SE vs. 74.7 % ± 0.9 SE; p = 0.011), however, the effect of BA is no longer significant (71.4 % ± 1.4 SE vs. 74.0 % ± 1.5 SE; p = 0.210). Based on these findings, BA has the strongest effect on overall performance between the ages of 12 to 13 years. Interestingly, the centre of this window is very close to the average menarche age (12.1±0.9SD years) of our participants.

In addition to the graphical analysis of Fig. 2, partial correlation and multiple regression analyses were also performed in the whole group in order to disentangle the *independent* effect of chronological age and biological age. We performed bootstrapping on 1000 samples to obtain confidence intervals for the partial correlation coefficients. The results are summarized in Tables 1 and 2 (results for the 11 subtests are summarized in Supplementary Tables 3 and 4).

**Table 1:** Partial correlations between chronological/biological age and WISC-IV overall performance, and performance on the four factors. Partial correlations between chronological age / biological age and WISC-IV broad abilities and overall performance. The correlations represent the *independent* effect of chronological age and biological age, with the effect of the other kind of age controlled for. 95CI = 95% confidence interval.

|  | Chronological Age | 95CI | p | Biological Age | 95CI | p |
| --- | --- | --- | --- | --- | --- | --- |
| Overall Performance | .295 | .130 -.449 | .001 | .328 | .142 -.488 | <.001 |
| Verbal Comprehension | .320 | .159 -.470 | <.001 | .060 | -.141-.252 | .524 |
| Perceptual Reasoning | .152 | -.040 -.343 | .104 | .075 | -.110-.262 | .425 |
| Working Memory | -.080 | -.272 -.118 | .395 | .360 | .201 -.504 | <.001 |
| Processing Speed | .173 | -.037 -.380 | .063 | .327 | .166 -.477 | <.001 |

$p < .01$
$p < .001$

**Table 2:** Multiple linear regressions for the effects of chronological/biological age on WISC-IV overall performance and on the four factors. Multiple linear regressions for chronological age / biological age on WISC-IV broad abilities and overall performance. B = unstandardized coefficients, beta = standardized coefficients, 95CI = 95% confidence interval.

|  |  | B | 95CI | beta | p | Model parameters |
| --- | --- | --- | --- | --- | --- | --- |
| Overall Performance | Chronological Age | .016 | .006 -.025 | .342 | .001 | F(2,114) = 49.444, p < .001<br>R = .682<br>R<sup>2</sup> = .465 |
|  | Biological Age | .016 | .007 -.024 | .386 | < .001 |  |
| Verbal Comprehension | Chronological Age | .034 | .015 -.052 | .443 | < .001 | F(2,114) = 19.433, p < .001<br>R = .504<br>R<sup>2</sup> = .254 |
|  | Biological Age | .005 | -.011 -.022 | .079 | .524 |  |
| Perceptual Reasoning | Chronological Age | .013 | -.003 -.030 | .222 | .104 | F(2,114) = 6.113, p < .01<br>R = .311<br>R<sup>2</sup> = .097 |
|  | Biological Age | .006 | -.009 -.021 | .108 | .425 |  |
| Working Memory | Chronological Age | -.008 | -.025 -.010 | -.108 | .395 | F(2,114) = 14.310, p < .001<br>R = .448<br>R<sup>2</sup> = .201 |
|  | Biological Age | .033 | .017 -.048 | .524 | < .001 |  |
| Processing Speed | Chronological Age | .014 | -.001 -.030 | .214 | .063 | F(2,114) = 31.733, p < .001<br>R = .598<br>R<sup>2</sup> = .358 |
|  | Biological Age | .026 | .012 -.039 | .421 | < .001 |  |

p < .01
p < .001

These analyses demonstrated that the independent effect of both BA and CA on overall performance is significant, substantial, and of very similar magnitude (pr=.298, p<.01 and .320, p<.001, respectively; B=.016, p<.01, and B=.016, p<.001, respectively). As discussed in the next section, this similar sized effect is not identical across specific cognitive abilities. On the contrary: the effect of BA and CA is strongly dissociable, only one or the other has a unique effect on specific abilities.

### Selective effects of biological and chronological age on specific cognitive abilities

Similarly to overall performance, regression and partial correlation analysis clearly dissociate the effect of maturity (BA) and experience (CA) on specific abilities reflected by the four indices of the WISC (Tables 1 and 2).

CA has a significant effect on Verbal Comprehension, independently of BA (pr=.320, p<.001; B=.034, p<.001), while BA does not have a significant effect independent of CA. For Working Memory and Processing Speed we identified the opposite pattern: BA has a significant effect independent of CA (pr=.360, p<.001 and .327, p<.001, respectively; B=.033, p<.001, and B=.026, p<.001, respectively).

A comparison of the partial correlations in Table 1 with the Pearson correlations in Supplementary Table 1 reveals that the independent effect of BA and CA do not manifest themselves in the latter. This should not come as a surprise given that BA and CA share most of their variance. Therefore, it is all the more remarkable that substantial independent effects could be statistically identified, and the effect of BA and CA could be dissociated for specific cognitive abilities.

Importantly, these effects are not driven by unique effects of individual subtests: as it is apparent from Supplementary Tables 3 & 4, the independent effect of BA and CA is uniform across each broad cognitive ability. That is, in each subtest that contributes to a given index score, the same significant independent effect was identified as in the index score itself that reflects a particular specific ability. This demonstrates the construct-level effect of BA and CA: they do not simply influence performance on individual subtests or narrow abilities, but rather on broad specific abilities.

There was no significant effect of BA or CA on Perceptual Reasoning.

### Age-specific maturity effects on specific cognitive abilities

Partial correlation and linear regression analyses both demonstrate that CA and BA selectively impact on different cognitive domains. Although overall performance on WISC-IV is determined both by CA and BA, the abilities reflected by the indices of the WISC-IV are either affected by CA or BA, suggesting that individual domains are either experience or maturation determined. However, as we have seen in Figs. 2b and 2c, CA and BA effects might depend on the actual age-range, and developmental windows may shift also for different cognitive abilities. In order to scrutinize the strong, age-specific selective effects of CA and BA that are suggested by Tables 1 and 2 in terms of age-specificity, we continued with the graphical analysis presented in Fig. 2a. The results for the Verbal Comprehension, Working Memory, Processing Speed and Perceptual Reasoning indices are shown in Fig. 3.

**Fig. 3:**
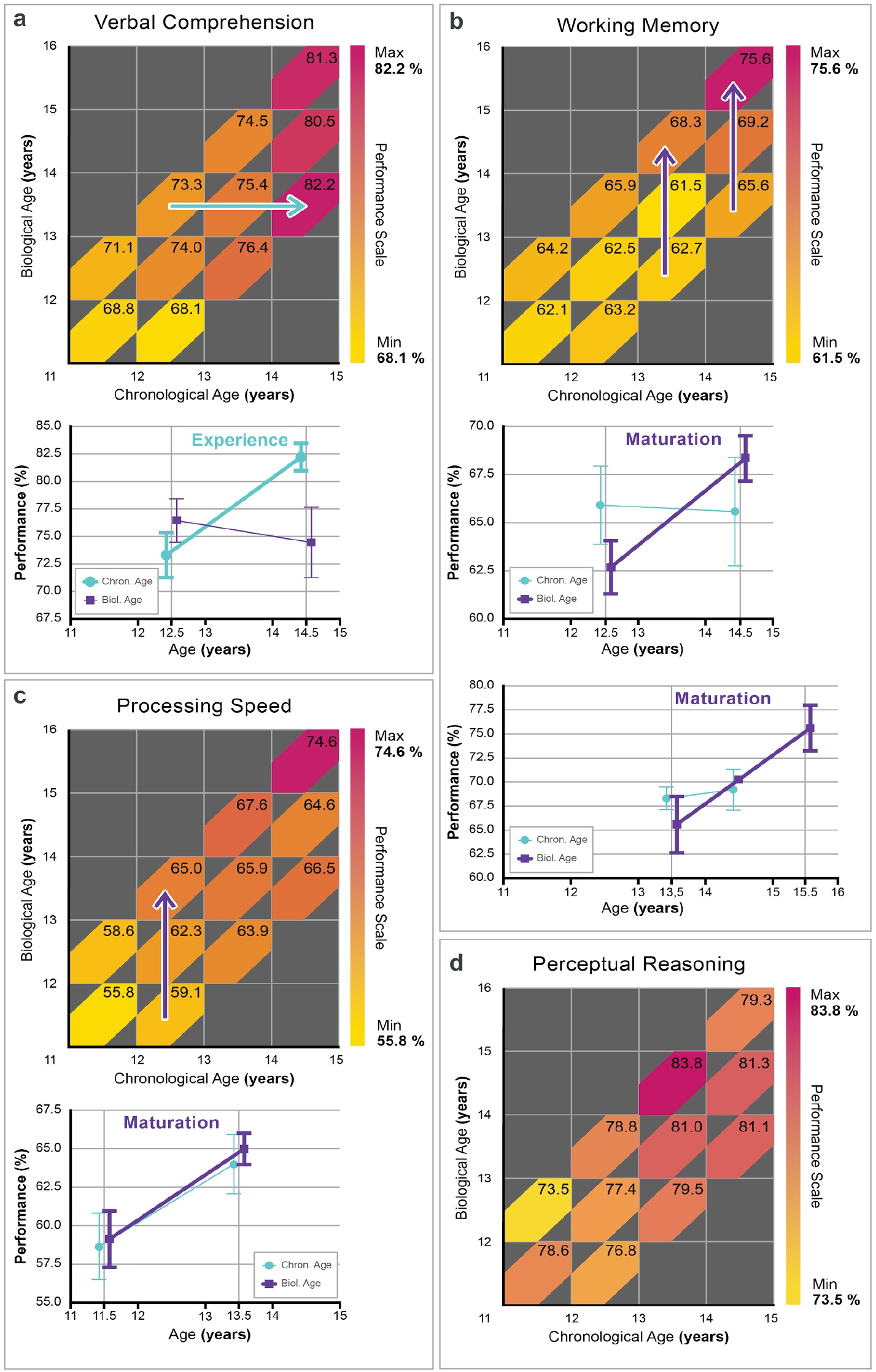
Performance in specific abilities as a function of biological and chronological age. Average performance (based on raw scores) for the four factors of WISC-IV was calculated as a % of maximum possible performance for each bin described in Fig.1b and expressed on a color scale (Performance scale). Higher performance is indicated by darker shades of red. Numbers within each colored bin express performance in % of maximum performance. → indicates bins where a significant biological age effect is present. → indicates bins where a significant chronological age effect is present. Line graphs show detailed performance within specific age-ranges where there are significant BA and/or CA effects according to one-way ANOVA (significant BA and/or CA effects are represented by bold lines). **a**, Verbal Comprehension is determined by chronological age, significant performance changes are only observed along the ‘Experience’ axis within the age-range of 12.5-14.5 years. **b**, Working Memory is determined by biological age, significant performance changes are only observed along the ‘Maturation’ axis within the age-ranges of 12.5-14.5 and 13.5-15.5 years. **c**, Processing Speed is determined by biological age, significant performance changes are only observed along the ‘Maturation’ axis within the age-range of 11.5-13.5 years, although the impact of experience can also be observed at a smaller extent. **d**, Perceptual Reasoning is not affected significantly by biological or chronological age within the investigated age-ranges.

Pearson correlation was calculated between each WISC index and BA and/or CA within each one-year wide BA/CA group. The correlations were Holm-Bonferroni corrected for multiple comparisons. We used one-way ANOVA to look for significant differences between the lower and upper bins within the particular BA/CA age-band with significant correlations. Coloured arrows in Fig. 3 indicate those BA/CA age-bands where significant differences were found.

Fig. 3a clearly shows that the Verbal Comprehension index of WISC-IV is determined by experience significantly at a constant 13.5 years of BA (73.3 % ± 1.3 SE vs. 82.2 % ± 1.2 SE; p < 0.001), while there is no significant maturity effect (76.4 % ± 2.0 SE vs. 74.5 % ± 3.2 SE; p = 0.591). Working Memory (Fig. 3b) on the other hand is BA dependent at a constant 13.5 and 14.5 years of CA (62.7 % ± 1.4 SE vs. 68.3 % ± 1.2 SE; p = 0.007 and 65.6 % ± 2.8 SE vs. 75.6 % ± 2.2 SE; p = 0.017, respectively), while there is no significant effect of experience (65.9 % ± 2.0 SE vs. 65.6 % ± 2.8 SE; p = 0.923 and 68.3 % ± 1.2 SE vs. 79.2 % ± 2.1 SE; p = 0.723, respectively). Processing Speed is showing a significant BA effect at a constant 12.5 CA (59.1 % ± 1.8 SE vs. 65.0 % ± 1.0 SE; p = 0.009), while the effect of experience does not reach significance (58.6 % ± 2.2 SE vs. 63.9 % ± 1.9 SE; p = 0.056, Fig. 3c). Perceptual Reasoning (Fig. 3d), corresponding to our analysis in Tables 1 and 2, does not seem to show any relevant BA or CA effects) within the investigated age-ranges.

## Discussion

We combined ultrasonic bone age estimation and a careful selection of participants according to their pubertal maturity levels to uncover the distinct effects of biological and chronological age on cognitive abilities. Unlike previous studies that employ less efficient control of pubertal maturity, our design allows for a clear-cut dissociation of the effects of maturity versus experience. By comparing the cognitive performance of participants with average, advanced and delayed skeletal maturity we revealed that overall cognitive performance is determined both by biological and chronological age. With respect to different broad cognitive abilities, we found that biological and chronological age selectively and independently influence them: among adolescents of the exact same chronological age, more mature individuals have higher memory capacity and processing speed, while among adolescents of the same maturity, those with higher chronological age have better verbal abilities. Our design also allowed for a detailed exploration of the shifting developmental windows of different cognitive abilities: we revealed age-specific effects with processing speed being affected by maturity levels at an earlier chronological age than working memory capacity.

The results have great importance from the perspective of fractionating cognitive abilities. Importantly, in this instance it is not simply an age effect dissociating cognitive abilities that is discovered, but the mechanism responsible (sexual maturation and the accompanied hormonal changes) is also identified. Relatedly, our results inform the debate about the adequate level of interpretation of IQ-test results in general^43,44^, and for clinical^45,46^ and school psychology^45,47–49^ in particular. Generally, there are three possible levels of the evaluation of performance in cognitive test batteries that consist of a number of subtests: 1) the level of subtests, 2) the specific factors, or 3) global IQ scores such as the Full Scale IQ (FSIQ) on the WISC.

Arguments in favour of interpreting only the global score usually rely on incremental validity analysis of predicting external criteria. Also, virtually all such studies employ factor analytic designs that prioritize the general factor (i.e., across-domain variance on cognitive tests), while residualizing broad ability factors. Both practices are problematic if there is no substantive reason to justify them^50,51^. In fact, we have suggested that without sufficient evidence to establish that variation in a single general mental ability causes individual differences in performance in cognitive ability tests, emphasis should be put on broad specific abilities instead of global indicators of performance^52^. Therefore, the theoretical background of test score interpretation is related to one of the oldest debates in psychology: is there a general mental ability that is measured by all kinds of cognitive ability tests regardless of their actual content? The theory of general intelligence (or *g*-theory) claims that there is, and practically states that psychometric *g,* a statistical representation of the across-domain variance between different tests, is the reflection of a psychological trait, general intelligence or general mental ability, and all tests measure this same trait to varying extent. However, the concept of general intelligence does not sit well with a number of findings from neuroscience and cognitive psychology. There have been recent accounts of the across-domain variance, as well as other main findings in intelligence research, that do not postulate a general intelligence, for instance mutualism^53,54^ or process overlap theory that we proposed^55,56^. The current results are in agreement with these latter theoretical accounts.

Historically, the WISC, similarly to the Wechsler Scales in general, were constructed under the strong influence of *g*-theory. Therefore, purportedly, “the subtests are different measures of intelligence, not measures of different kinds of intelligence”^57^ (p. 64). More recent versions of the Wechsler Scales saw the addition of four broad ability indices besides Full Scale IQ scores, but the emphasis is still often on FSIQ as a proxy for *g*. Yet our results highlight the importance of interpreting test results at the level of broad specific abilities, too. The history of intelligence test interpretation saw the influence of theories of cognitive abilities only recently, in what is usually called the 4th and currently last wave^58,59^. The theoretical implications of the finding that different broad abilities are affected by different causal factors during development should indeed be informative for both test users and test constructors.

Our findings can also inform research on the causes of IQ changes in adolescence. A child’s relative standing in their peer group – reflected by IQ, an age-standardized indicator of cognitive abilities – can undergo substantial change. It has been demonstrated that such changes are accompanied by corresponding changes in the structure and organization of the brain^60,61^, therefore they are genuine shifts in one’s standing on the peer ability ranking and not reflections of measurement error. Our results indicate that the ultimate cause of a teen outdriving their peers or, on the contrary, falling behind in one’s age group, might be individual differences in biological maturation, driving the acceleration of the development of abilities –similarly to how children can move substantially ahead or behind in the height order of their class as the result of variation in the timing of maturation.

Our results demonstrate that individual differences in maturation in adolescents who are of the same age have a selective effect on some broad abilities, independently of chronological age. This finding is enormously important for practical issues in school psychology. For instance, differences in school performance between children in the same class might in part be due to differences in hormonal maturation. Additionally, this result is also informative for planning intervention policies, since children who are delayed in their hormonal development will eventually “catch up.” Therefore, the institutional measurement of bone age might be warranted for children who exhibit a pattern of developmental lags one would expect on the basis of a shortfall of hormonal maturation compared to peers.

With respect to the age-specific effects of maturity on different cognitive abilities, Working Memory and Processing Speed Indices of the WISC-IV stand out in our study. In addition to both being strongly correlated with the level of physical maturity, processing speed is affected by biological age earlier (12-13 yrs., Fig. 3c) than working memory (13-15 yrs., Fig. 3b). Although a strong association between processing speed and working memory during development has been asserted^62,63^, the picture is confounded unless controlling for age and maturity independently. The largest available population study including these variables is the Adolescent Brain Cognitive Development (ABCD) Study (https://abcdstudy.org). At its current stage, ABCD provides a conclusive analysis on mostly prepubertal 10-year-old children demonstrating that processing speed and working memory are not related at this age^17^. Our results corroborate this finding revealing the difference in the maturational timing of the two behavioural measures, and also support an interpretation emphasizing their diverse neural background. While processing speed is generally associated with white matter volume^64,65^, working memory is more specifically related to the late maturing or frontoparietal core network^66–69^.

Although working memory (Fig. 3b) seems to fulfil our predictions with respect to an exclusively maturation-determined (Fig. 1c) developmental process, verbal comprehension (Fig. 3a) exhibits a uniquely experience-determined (Fig. 1c) process. Our results on the Verbal Comprehension Index (Fig. 3a) and the related subtests (Supplementary tables 3 and 4) are in agreement with previous results which indicate that the verbal skills the WISC measures are determined by age and schooling. For instance, unlike nonverbal abilities, performance on the verbal subtests keeps increasing in adulthood and peaks in middle to old age – in fact, of all the WISC subtests these are the ones to peak latest in life^35^. Additionally, previous studies that employed a methodology capitalizing on a possible discrepancy between a child’s chronological age and their school grade in order to dissociate the effect of schooling from other kinds of experience that improve with age, found that in children schooling has a particularly strong effect on individual differences in this ability^37^. Therefore, our conclusion that chronological age has a selective effect on the Verbal Comprehension Index, independent of maturity, is in accord with previous findings. The extent to which this effect is due to schooling in particular warrants further research.

In addition to the effect of chronological age, the lack of an effect of maturation is also important. While one could plausibly expect such an effect provided the cognitive basis of language development, the Verbal Comprehension Index of the WISC, despite its name, does not strongly tap linguistic cognition. In fact, the same factor has been interpreted as Gc (Comprehension-knowledge) in the CHC model^70,71^, which is defined as cultural knowledge acquired during education and life experiences^33^. Similarly, in forerunners of the CHC model the same construct (Gc) was called Crystallized intelligence and Acculturation knowledge^72,73^. From this perspective a lack of biological influence is to be expected; in fact, our results contribute to the construct validity of the above conceptualization of Gc.

The Perceptual Reasoning Index of WISC-IV (Fig. 3d) does not seem to show any relevant biological or chronological age effects within the investigated age-ranges. This is somewhat against expectations since perceptual development extends well into the teenage years as we have demonstrated earlier^74,75^. While lower-level perceptual functions are adult-like in the early years^76^, integrative processes of perceptual organization mature relatively late^74,75^. On the other hand, the canonical back to front progression of brain maturation^77–79^ with the corresponding functional developmental trajectories^8,80,81^ may suggest that the developmental window (Fig. 1c) for the perceptual functions tapped by the probes in the Perceptual Reasoning Index is at an earlier age than our investigated range. Another possibility is that the probes within this Index are less reliable measures of function than those within the other scales. Detailed psychophysical studies employing the suggested biological/chronological age dissociation paradigm might decide between these options. Although we have already dissociated the role of maturity and experience in early visual development^82^, it would be extremely important to do the same beyond childhood since later perceptual alterations might be at the core of behavioural dysfunction in several neurodevelopmental disorders^83–87^.

The findings presented here have great relevance for clinical neuroscience, especially with respect to the so-called “connectopathies” such as schizophrenia or autism. Puberty related hormonal influence on the excess elimination of cortical synapses (“pruning”) have been indicated to play a role in early onset schizophrenia with evidence for overpruning^88,89^, while adolescent underpruning has been indicated among the various age-specific anatomical alterations in autism^90–93^. Variation in puberty onset times seems to be related to eating disorders and depression, with early maturing adolescents having a greater risk to develop those^94,95^. Clearly, adolescence is an incredibly sensitive period of hormone-dependent brain organization. Since the manipulation of gonadal steroid levels is not an option in human research, direct evidence is generally difficult to obtain in this field. Therefore, the establishment of a reliable pubertal maturity indicator, and an associated neurotypical baseline of brain structure and function is definitely necessary for a better understanding of the pubertal remodelling of the human brain that might lead to efficient clinical interventions.

In general, human neurocognitive development has a uniquely protracted course and the details on factors that determine maturational trajectories of brain connectivity are only beginning to emerge^96^. The orchestrating function of pubertal hormones behind overall adolescent progress seems clear and assumes an explanatory role. Pubertal hormone levels are correlated with bone age^26–28^, time bodily growth^97^, cortical pruning^98,99^, and emotional^1,4^ and cognitive^100,101^ development in adolescence. The cascade of events during the pubertal transformation of a child into an adult is coordinated on individual timescales, and any attempt to assess adolescent brain and cognitive function requires the determination of the individual timescales. Emphasizing the importance of individual differences in neurodevelopment^1,12^ and parsing the genetic versus environmental influences on the human brain^102^ are among those pertinent ideas that should move the field forward. However, it is unlikely that a nuanced account of morphological and functional neurodevelopment will be available without properly addressing the trade-off between biological maturity and life-time experience. The work presented here seems to fill this gap. It provides a suitable paradigm to address the mentioned trade-off, and thereby establishes a methodological niche for future studies on adolescent brain development.

It seems that bone-age assessment has proven to provide a much better grasp on puberty stages than previous attempts and helped us to show the direct relationship between cognitive development and physical maturity. The implications for brain research are clear: it is essential to determine what the underlying cortical and subcortical structures are behind the maturity dependent cognitive functions we found, and to reveal the maturational changes in the human brain initiated and determined by the onset of puberty. This would shed light on yet uncovered developmental mechanisms and help us to draw developmental trajectories at a very fine scale.

The current project is constrained to study adolescent girls only which might appear as a limitation. However, it is unavoidable to make a choice if the primary purpose is to clearly dissociate the effects of maturation and experience on cognitive development. Involving both sexes would have confounded the clear-cut paradigm we developed for the dissociation (Fig. 1a). Since male pubertal onset is delayed by 1-1.5 years with respect to that of females^39–41^, results from girls and boys could not have been analysed together. We chose to include girls because menarche age serves as valuable additional information confirming the validity of bone age assessments. Based on the conclusive results with girls that indicate a strong role of biological age in cognitive development one might assume that a similar effect will also be observed in boys. Indeed, we would expect that maturity is a relevant factor in the general cognitive development of boys as well, however, due to the fact that female and male developmental trajectories are determined by very different gonadal hormone levels^4,103^, we also anticipate alternative trajectories in terms of specific cognitive abilities. Such trajectories have recently been indicated, e.g., in visuospatial processing^104^ and visual decision making^105^.

Another restriction we felt important to introduce is to only involve students at top-level high schools. Since we primarily sought to study the effect of maturity on adolescent development, large differences in socio-economic or socio-cultural backgrounds as well as the preselection processes introduced by schools would have confounded our data. We chose to include schools at the high end since entrance exam scores of accepted students vary the least in those. Based on the findings of the current study, we would expect to find meaningful discrepancies in cohorts with socio-economic or socio-cultural handicaps where sexual precocity is more frequent^3,39^.

In spite of the limitations mentioned above, we successfully dissociated the influence of chronological and biological age on a wide range of cognitive abilities and predicted that different cognitive abilities will be relying on maturity and experience at different rates (Fig. 1c). This prediction is confirmed by the results. Although it may have been anticipated that time and experience boost development, the finding that biological maturation assessed through skeletal maturation has such a profound effect on the development of cognitive abilities is striking.

Considering the remarkable association between skeletal maturity and cognitive ability unearthed here, ultrasonic bone age assessment, together with the paradigm allowing for the clear dissociation of maturity and experience, seems to be a valuable asset for research purposes. It is indeed timely to introduce an accurate, objective, non-invasive and ethically impeccable pubertal maturity measurement technique into adolescent research in order to accurately estimate adolescent developmental trajectories. In addition to the interpretable findings, we hope that our methodological invention will contribute to adolescent brain and cognitive development research, and to the development of prevention and intervention techniques in the related clinical fields.

In conclusion, we have introduced bone age assessment into adolescent research to replace less reliable methods of pubertal maturity estimation. This method allowed us to disentangle the independent effects of chronological and biological age on adolescent cognitive abilities. Thereby, we compared cognitive performance of participants with average, advanced and delayed skeletal maturity and uncovered the striking impact of biological age on overall cognitive performance in addition to the known and expected effect of chronological age. With respect to the tested broad cognitive abilities, we discovered that more mature individuals within the same chronological age group have higher working memory capacity and processing speed, while those with higher chronological age have better verbal abilities, independently of their maturity levels. The paradigm introduced here is expected to fill a methodological niche for future studies on adolescent brain development, and the results already obtained will advance developmental neuroscience, school and clinical psychology as well as the study of the nature of intelligence and the determinants of individual differences in cognitive abilities.

## Materials and Methods

### Participants

Due to the fact that median age at menarche is between 12.5-13.5 years of age in females^40,106^, and male pubertal onset lags behind by about 1-1.5 years^39–41^, it would have been difficult to include both sexes in the current study. Since our aim is to investigate whether biological age has a significant effect on cognitive development as a function of maturity, the delayed maturation of boys would have confounded the design described in Fig. 1a. Therefore, we only tested female participants (mean menarche age of the whole sample is 12.1±0.9SD years).

We recruited participants mainly by contacting schools and advertising online. Our participants were selected from high schools that belong to the top 10% in Hungary, according to a ranking based on university enrolment. Since Hungarian high schools accept pupils based on an entrance exam, and the highly ranked schools select students with the highest scores, it appeared relevant to limit the range of participating schools to avoid the confounding effect of school ranking. The top 10% seemed a good choice because these schools accept students uniformly with scores varying only slightly around maximal performance, while performance varies much more in high-average or average schools.

Parents filled out a questionnaire on demographic data, parental education, height, age at birth, number of siblings, residential area, time of menarche, handedness, fine motor function, musical training, and visus. Parents also reported any history of developmental disorders, learning disability, neurological disorders, sleep disorder, attention deficit disorder or recent injury to the wrist area on the non-dominant hand. There was only one parent who reported attention deficit disorder in their child who was then excluded from further testing.

All participants went through an ultrasonic bone age screening procedure (see Procedure). Following the screening, we selected an approximately equal number of participants into the BA/CA defined bins as shown in Fig. 1b. In order to form discrete, one-year-wide bins without any chance of overlap between the average, advanced and delayed categories, we defined four CA groups (Table 3), and constrained bin sizes in terms of BA: an average maturity bin is defined as CA-0.5≤BA≤CA+0.5 years; an advanced maturity bin is defined as CA+1.5≥BA>CA+0.5; a delayed maturity bin is defined as CA-1.5≤BA<CA-0.5.

**Table 3:** Descriptive statistics of participants’ age. Descriptive statistics of the selected 117 participant cohort in terms of CA and BA. Maturity levels are color coded according to Fig 1b. Each CA group starts at the birthday and ends just before the birthday of a participant (e.g.: 11.00-11.99 year). BA determines bin-sizes in a similar manner. The exact number of participants in each bin is presented in Fig. 1b.

| CA groups | Maturity level | CA Mean (SD) | BA Mean (SD) |
| --- | --- | --- | --- |
| 11-12 years | Advanced | 11.48 (.22) | 12.45 (.21) |
|  | Average | 11.63 (.17) | 11.60 (.26) |
| 12-13 years | Advanced | 12.53 (.25) | 13.47 (.24) |
|  | Average | 12.48 (.28) | 12.52 (.24) |
|  | Delayed | 12.37 (.28) | 11.54 (.22) |
| 13-14 years | Advanced | 13.68 (.33) | 14.53 (.20) |
|  | Average | 13.50 (.30) | 13.47 (.29) |
|  | Delayed | 13.49 (.23) | 12.53 (.29) |
| 14-15 years | Advanced | 14.48 (.32) | 15.68 (.35) |
|  | Average | 14.36 (.24) | 14.40 (.28) |
|  | Delayed | 14.51 (.23) | 13.55 (.31) |

Power calculations for the linear regression analysis indicated that a sample size near N=120 was warranted in order to avoid type II error. In a pilot BA assessment study, we found that the percentage of adolescents with advanced or delayed maturation is around 20% in each group. Accordingly, we determined that the total number of participants to be invited for BA screening should be 300, with each CA group including 75 screened participants. With 300 participants screened, we predicted that there will be at least 10 participants within each bin. Therefore, 300 female students between the ages of 11 and 15 years were invited for BA screening. In order to eliminate the prospect of endocrinological complications, we excluded extremely advanced (BA>CA+1.5, 21 girls, 7.0%) and extremely delayed (BA<CA-1.5, 16 girls, 5.3%) participants from further testing. An additional 34 (11.3 %) participants did not fit into any of our predetermined bins (falling just outside of the hexagons in Fig. 1b). 105 (35.0%) participants fell into average, 67 (22.3%) participants into advanced, and 57 (19.0%) participants into delayed bins of skeletal maturation. Out of the 229 participants falling into the predetermined bins, 53 girls have not accepted our invitation for further testing, and 59 girls were not invited back since we reached our predetermined sample size.

The final cohort described in Fig. 1b and in Table 3 consisted of 117 girls. These 117 girls participated in WISC-IV testing. All of them speak Hungarian as their mother tongue. 17 participants are left-handed and there is a mixed-handed participant. One of the participants got distracted during the administration of the Cancellation subtest, therefore their result on this subtest was discarded from the analysis.

The Hungarian United Ethical Review Committee for Research in Psychology (EPKEB) approved the study (reference number 2017/84). Written informed consent was obtained from all subjects and their parents. Participants were given gift cards (approx. EUR 15 value each) for their attendance.

### Procedure

#### Bone age assessment

After body height (DKSH anthropometer, Switzerland Ltd, Zurich, Switzerland) and body weight (Seca digital scale) measurements, we assessed skeletal maturity (bone age) with an ultrasonic device (Sunlight BonAge, Sunlight Medical Ltd, Tel Aviv, Israel). Ultrasonic testing prevents children from being exposed to radiation in contrast to earlier, X-ray based methods, and at the same time, ultrasonic estimations highly correlate with X-ray estimations^30^. These features make ultrasonic screening optimal for research purposes. Ultrasonic bone age estimation is based on the changing acoustic conductivity of forearm bones during growth.

Ultrasonic bone age estimation was carried out at the school of the participants or at the Research Centre for Sport Physiology at the University of Physical Education, Budapest.

We performed measurements on the left hand and wrist region. Participants placed their arm on the armrest surface between the transducers. We adjusted the transducers to the forearm growth zone (to the connection of the distal epiphysis and diaphysis). Then, at the initial position, the device adhered to the forearm at a pressure of approximately 500 g and emitted ultrasound at a frequency of 750 kHz to the measurement site for each measurement cycle. One measurement cycle lasted about 20 seconds and was repeated five times. Between the measurement cycles, the transducers rose up two millimetres in the palm-back direction. The device estimates bone age (in years and months) by measuring the speed of sound (SOS) and the distance between the transducers, using algorithms based on gender and ethnicity^107^. The same person performed all measurements.

### WISC-IV testing

Within a maximum of 3 months after the bone-age assessment of each participant we administered 11 subtests of the Hungarian adaptation of the Wechsler Intelligence Scale for Children IV - WISC-IV^36^: Verbal Comprehension Index - Similarities, Vocabulary, Comprehension; Perceptual Reasoning Index – Block Design, Visual Puzzles, Matrix Reasoning; Working Memory Index – Digit Span, Letter-Number Sequencing; Processing Speed Index – Coding, Symbol Search, Cancellation. For the justification of subtest selection in the current study please see Supplementary information/WISC subtest selection. The administration of each subtest followed the standard testing and scoring procedure according to the test manual^108^.

WISC-IV subtests were administered in one session and scored according to the manual. Four trained school or clinical psychologists carried out the testing at the Laboratory for Psychological Research at PPCU. Scoring of the verbal comprehension subtests have subjective elements, therefore, we assessed interscorer reliability. Three cases were randomly selected from the sample and scored independently by all four psychologists. Intraclass correlation coefficient of interscorer reliability was 0.992.

## Acknowledgements

The authors thank T. Jáger for generating the figures and tables. We also thank the generous and amazing parents, adolescents and schools who participated in this project, and those trained psychologists who carried out WISC-IV testing. The project was supported by the National Research, Development and Innovation Office of Hungary (Grant K-134370 to I.K., Grant PD-17-125360 and Grant KH-18-130424 to K.K.) and by the Eötvös Loránd Research Network, Hungary (ELRN-PPCU Adolescent Development Research Group). The funders had no role in study design, data collection and analysis, decision to publish or preparation of the manuscript.

## Author Contributions

I.K. and F.G. conceptualized and designed the paradigm; G.O. contacted schools, organized and carried out subject screening, preselection of subjects and data collection; K.U. and Z.T. carried out the anthropometric measurements and data analysis; F.G. and K.K. carried out the statistical analysis; G.P., A.B. and H.S. conducted analysis and contributed to data interpretation; I.K. supervised the entire work. I.K. and K.K. drafted the manuscript. All authors discussed the results, contributed to the text, and approved the final version of the manuscript for submission.

## Supplementary Information

### WISC subtest selection

We purported to measure each broad ability with three tests. In the case of Verbal Comprehension and Perceptual Reasoning the ten core tests of the battery already include three tests of each. For Perceptual Speed, there are two core tests in the standard battery, and they can be supplemented with a third test, Coding, which we also administered. In the case of Working Memory there are two core tests in the standard battery and a third test, Arithmetic, is available as supplementary.

However, the status of the Arithmetic subtest is controversial. Upon examining the content of the test, it appears as a complex measure that taps on various domains at the same time: quantitative knowledge, quantitative reasoning, working memory, and even verbal comprehension. In fact, the Cattell-Horn-Carroll (CHC) model recognizes two different aspects of individual differences in math-related cognition: the separate broad ability factor ‘Quantitative knowledge’ reflects acquired quantitative or numerical knowledge, but not reasoning with such knowledge, while ‘Quantitative reasoning’, a narrow factor under the broad ability factor ‘Fluid reasoning’ represents reasoning with numerical material^33^.

In order for it to be a better measure of Working memory, the publishers of the WISC-IV have substantially modified the Arithmetic test from previous versions: the math-knowledge load was reduced, and the working memory demands were increased^109^. Despite this, several studies investigating the factor structure of the WISC-IV found that it is still not a pure measure of working memory; it has been found that besides memory, it measures fluid reasoning^70,110^ and/or crystallized intelligence/comprehension & knowledge^71,111^, too. Even a study that confirmed Arithmetic as a measure of Working memory found that its factor loading is much smaller than of the core working memory subtests^112^.

A study fitted a model in which the Arithmetic test was removed from the 4-factor structure of the WISC and was directly measuring the higher order *g* factor^113^. Such a model is equivalent to one in which a separate factor is added, of which Arithmetic is the single indicator. Therefore, such a factor is statistically redundant, but substantively it is more plausible to claim that Arithmetic is a measure of a fifth broad ability than a direct measure of *g*. Indeed, it appears that the main difficulty from a latent variable modelling perspective is that Arithmetic might be the single indicator of a fifth broad ability, only allowing for suboptimal models.

The manual did in fact consider such a 5-factor solution but discarded it because it did not improve model fit over the 4-factor solution. Yet, importantly from our perspective, the 5-factor solution with Arithmetic as the only indicator of the fifth factor did improve model fit in a particular age group: in 11–16-year-old children^109^. Since this is the exact age range we were targeting, after considering all the above evidence we decided against administering the Arithmetic test as a supplementary test of Working Memory.

---

**Table S1:** Descriptive statistics. Descriptive statistics of WISC-IV broad abilities and overall performance (upper panel), and individual subtests. M = mean, SD = standard deviation (lower panel). VC = Verbal Comprehension, PR = Perceptual Reasoning, WM = Working Memory, PS = Processing Speed

|  | M | SD | Range | N |
| --- | --- | --- | --- | --- |
| Overall Performance | 71.34 | 4.89 | 60.15 - 80.84 | 117 |
| Verbal Comprehension | 75.01 | 8.13 | 53.24 - 90.02 | 117 |
| Perceptual Reasoning | 79.18 | 6.48 | 60.99 - 91.89 | 117 |
| Working Memory | 65.29 | 7.41 | 48.54 - 84.06 | 117 |
| Processing Speed | 63.81 | 7.22 | 43.94 - 78.37 | 117 |
| VC – Similarities | 72.44 | 8.94 | 50.00 - 90.91 | 117 |
| VC – Vocabulary | 81.77 | 9.69 | 57.35 - 98.53 | 117 |
| VC – Comprehension | 70.82 | 9.39 | 50.00 - 90.48 | 117 |
| PR – Block Design | 80.74 | 11.39 | 44.12 - 98.53 | 117 |
| PR – Visual Puzzles | 74.88 | 8.86 | 57.14 - 100.0 | 117 |
| PR – Matrix Reasoning | 81.93 | 8.60 | 57.14 - 97.14 | 117 |
| WM – Digit Span | 60.12 | 10.06 | 37.50 - 87.50 | 117 |
| WM – Letter -Number Sequencing | 70.46 | 6.86 | 53.33 - 90.00 | 117 |
| PS – Coding | 54.30 | 9.56 | 29.41 - 78.15 | 117 |
| PS – Symbol Search | 55.88 | 8.66 | 33.33 - 75.00 | 117 |
| PS – Cancellation | 81.39 | 9.83 | 50.74 - 97.06 | 116 |

**Table S2:** Correlations for the four factors of WISC-IV. Correlations between chronological age / biological age and WISC-IV broad abilities and overall performance. 95CI = 95% confidence interval

|  | Chronological Age | 95CI | p | Biological Age | 95CI | p |
| --- | --- | --- | --- | --- | --- | --- |
| Overall Performance | .627 | .514 - .726 | <.001 | .643 | .533 - .739 | <.001 |
| Verbal Comprehension | .495 | .343 - .643 | <.001 | .410 | .246 - .573 | <.001 |
| Perceptual Reasoning | .306 | .124 - .472 | .001 | .275 | .105 - .432 | .003 |
| Working Memory | .271 | .091 - .440 | .003 | .441 | .310 - .558 | <.001 |
| Processing Speed | .534 | .380 - .665 | <.001 | .582 | .443 - .694 | <.001 |

p < .01
p < .001

**Table S3:** Partial correlations for the subtests of WISC-IV. Partial correlations between chronological age / biological age and WISC-IV subtests. The correlations represent the *independent* effect of chronological age and biological age, with the effect of the other kind of age controlled for. 95CI = 95% confidence interval. VC = Verbal Comprehension, PR = Perceptual Reasoning, WM = Working Memory, PS = Processing Speed

|  | Chronological Age | 95CI | p | Biological Age | 95CI | p |
| --- | --- | --- | --- | --- | --- | --- |
| VC – Similarities | **.209** | **.039 -.370** | **.025** | .077 | -.094 -.248 | .415 |
| VC – Vocabulary | **.246** | **.083 -.399** | **.008** | .067 | -.125 -.258 | .479 |
| VC – Comprehension | **.336** | **.182 -.488** | **<.001** | .016 | -.171 -.200 | .867 |
| PR – Block Design | .103 | -.090 -.323 | .272 | .140 | -.062 -.328 | .136 |
| PR – Visual Puzzles | .109 | -.073 -.272 | .246 | -.097 | -.277 -.091 | .301 |
| PR – Matrix Reasoning | .100 | -.059 -.273 | .289 | .079 | -.109 -.265 | .399 |
| WM – Digit Span | -.127 | -.313 -.075 | .177 | **.362** | **.210 -.500** | **<.001** |
| WM – Letter - Number Sequencing | -.035 | -.216 -.153 | .707 | **.270** | **.095 -.432** | **.004** |
| PS – Coding | .116 | -.053 -.289 | .216 | **.288** | **.142 -.419** | **.002** |
| PS – Symbol Search | .170 | -.033 -.367 | .070 | **.208** | **.021 -.386** | **.025** |
| PS – Cancellation | .100 | -.090 -.303 | .290 | **.201** | **.049 -.351** | **.032** |

p < .05
p < .01
p < .001

**Table S4:**
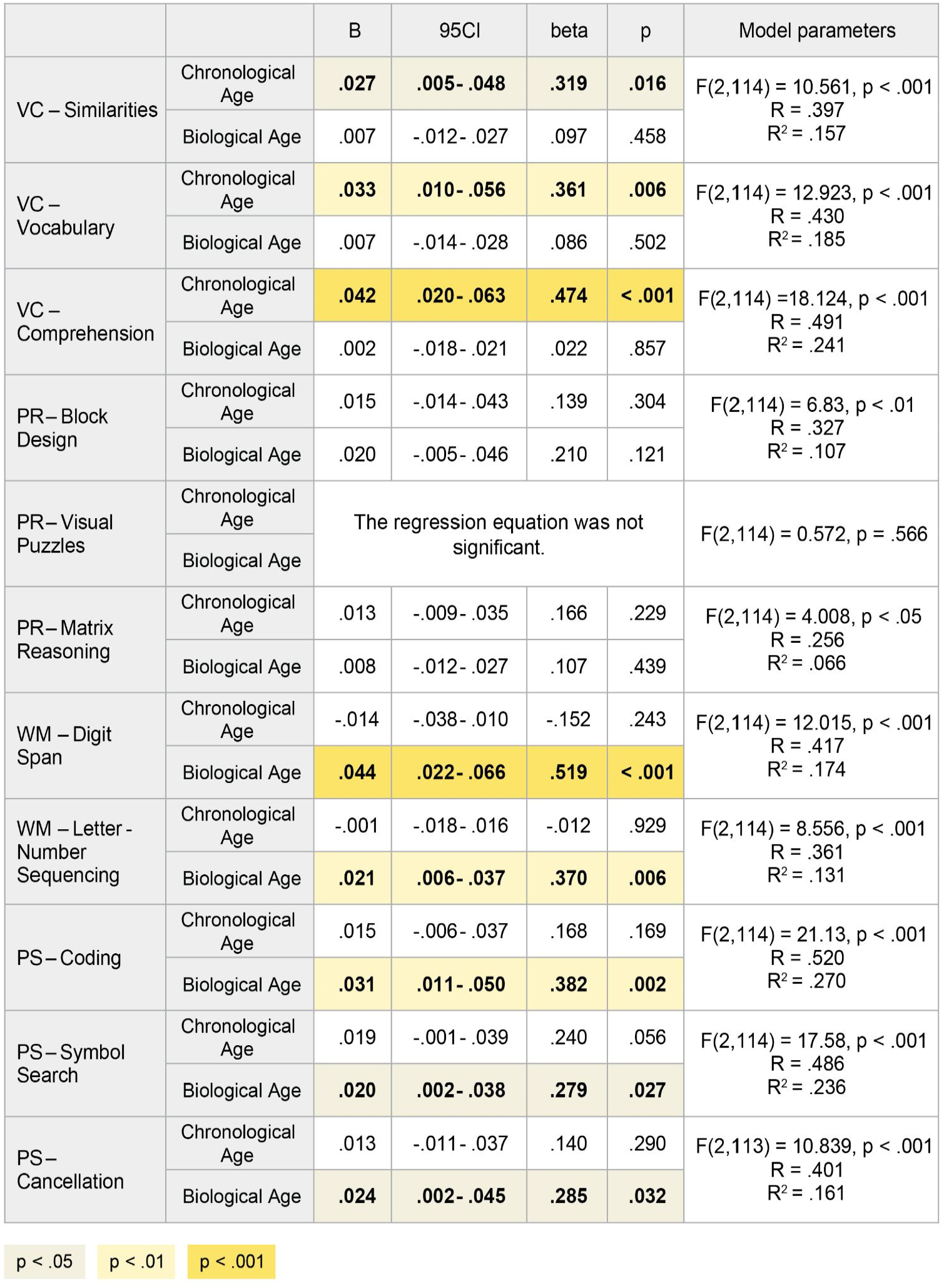
Linear regressions for the subtests of WISC-IV. Multiple linear regressions for chronological age / biological age on WISC-IV broad abilities and overall performance, B = unstandardized coefficients, beta = standardized coefficients, 95CI = 95% confidence interval.VC = Verbal Comprehension, PR = Perceptual Reasoning, WM = Working Memory, PS = Processing Speed.

|  |  | B | 95CI | beta | p | Model parameters |
| --- | --- | --- | --- | --- | --- | --- |
| VC – Similarities | Chronological Age | **.027** | **.005- .048** | **.319** | **.016** | F(2,114) = 10.561, p < .001<br>R = .397<br>R<sup>2</sup> = .157 |
|  | Biological Age | .007 | -.012- .027 | .097 | .458 |  |
| VC – Vocabulary | Chronological Age | **.033** | **.010- .056** | **.361** | **.006** | F(2,114) = 12.923, p < .001<br>R = .430<br>R<sup>2</sup> = .185 |
|  | Biological Age | .007 | -.014- .028 | .086 | .502 |  |
| VC – Comprehension | Chronological Age | **.042** | **.020- .063** | **.474** | **< .001** | F(2,114) = 18.124, p < .001<br>R = .491<br>R<sup>2</sup> = .241 |
|  | Biological Age | .002 | -.018- .021 | .022 | .857 |  |
| PR – Block Design | Chronological Age | .015 | -.014- .043 | .139 | .304 | F(2,114) = 6.83, p < .01<br>R = .327<br>R<sup>2</sup> = .107 |
|  | Biological Age | .020 | -.005- .046 | .210 | .121 |  |
| PR – Visual Puzzles | Chronological Age | The regression equation was not significant. |  |  |  | F(2,114) = 0.572, p = .566 |
|  | Biological Age |  |  |  |  |  |
| PR – Matrix Reasoning | Chronological Age | .013 | -.009- .035 | .166 | .229 | F(2,114) = 4.008, p < .05<br>R = .256<br>R<sup>2</sup> = .066 |
|  | Biological Age | .008 | -.012- .027 | .107 | .439 |  |
| WM – Digit Span | Chronological Age | -.014 | -.038- .010 | -.152 | .243 | F(2,114) = 12.015, p < .001<br>R = .417<br>R<sup>2</sup> = .174 |
|  | Biological Age | **.044** | **.022- .066** | **.519** | **< .001** |  |
| WM – Letter - Number Sequencing | Chronological Age | -.001 | -.018- .016 | -.012 | .929 | F(2,114) = 8.556, p < .001<br>R = .361<br>R<sup>2</sup> = .131 |
|  | Biological Age | **.021** | **.006- .037** | **.370** | **.006** |  |
| PS – Coding | Chronological Age | .015 | -.006- .037 | .168 | .169 | F(2,114) = 21.13, p < .001<br>R = .520<br>R<sup>2</sup> = .270 |
|  | Biological Age | **.031** | **.011- .050** | **.382** | **.002** |  |
| PS – Symbol Search | Chronological Age | .019 | -.001- .039 | .240 | .056 | F(2,114) = 17.58, p < .001<br>R = .486<br>R<sup>2</sup> = .236 |
|  | Biological Age | **.020** | **.002- .038** | **.279** | **.027** |  |
| PS – Cancellation | Chronological Age | .013 | -.011- .037 | .140 | .290 | F(2,113) = 10.839, p < .001<br>R = .401<br>R<sup>2</sup> = .161 |
|  | Biological Age | **.024** | **.002- .045** | **.285** | **.032** |  |

p < .05 p < .01 p < .001

